# Loss of 20-HETE-GPR75 Signaling Impairs Renal Hemodynamic Adaptation in Sepsis

**DOI:** 10.64898/2026.09.14.751621

**Authors:** Yaqi Han, Huaxing Zhang, Leying Li, Lixia Liu, Yanling Yin, Yuhong Chen, Xin Lin, Ruike Yang, Congcong Zhao, ZhenJie Hu

## Abstract

**BACKGROUD:** Sepsis is a life-threatening condition characterized by a dysregulated host response to infection. It frequently precipitates sepsis-associated acute kidney injury (SA-AKI). 20-Hydroxyeicosatetraenoic acid (20-HETE), is an important regulator of vascular tone. We aim to investigate whether disruption of 20-HETE signaling contributes to septic renal dysfunction.

**METHODS:** In the animal study, Sprague-Dawley rats underwent the cecal ligation and puncture to induce sepsis and SA-AKI. Comprehensive cardiac and renal ultrasonographic assessments were performed in all animals. Blood samples were collected for biochemical assessment of renal function and quantification of circulating 20-HETE levels. Kidney tissues were harvested for analysis of the 20-HETE–GPR75–PLC/PKC signaling pathway and histological evaluation by hematoxylin and eosin staining. In the clinical study, circulating 20-HETE levels and renal hemodynamic parameters derived from point-of-care ultrasound were evaluated in healthy individuals and patients with sepsis. Venous blood samples were collected for quantification of serum 20-HETE concentrations.

**RESULTS:** Firstly, CLP-induced septic rats produced a biphasic hemodynamic response, with early compensation followed by late cardiovascular decompensation, accompanied by progressive renal dysfunction, tubular injury, depletion of circulating and renal 20-HETE, and reduced renal GPR75 expression. Secondly, Restoration of 20-HETE signaling with a pharmacological analog improved systemic and renal hemodynamics, increased MAP, reduced serum creatinine, blood urea nitrogen, and ameliorated renal pathological injury. These protective effects were partially attenuated by PKC inhibition, implicating PKC-dependent signaling in 20-HETE-mediated renal protection. Moreover, in septic patients, circulating 20-HETE concentrations were reduced compared with healthy individuals and were lowest among those with SA-AKI, paralleling alterations in renal hemodynamic parameters measured by point-of-care ultrasound.

**CONCLUSIONS:** Sepsis-associated depletion of 20-HETE accompanies impaired systemic and renal hemodynamic regulation and is associated with the development of SA-AKI. Restoration of 20-HETE signaling improves hemodynamic function and attenuates renal injury, potentially involving GPR75-associated PLC/PKC signaling.

## INTRODUCTION

Sepsis is defined as life-threatening organ dysfunction caused by a dysregulated host response to infection. ^1^Despite advances in critical care medicine, sepsis remains a leading cause of acute kidney injury (AKI) among critically ill patients, accounting for a substantial proportion of AKI cases in the intensive care unit (ICU).^2^ When AKI develops in the setting of sepsis, it is referred to as sepsis-associated acute kidney injury (SA-AKI).^3,4^ Recently, the Acute Disease Quality Initiative (ADQI) 28 Workgroup established a consensus definition for SA-AKI, requiring the coexistence of sepsis according to the Sepsis-3 criteria and AKI according to Kidney Disease: Improving Global Outcomes (KDIGO) criteria, with AKI occurring within 7 days after sepsis diagnosis.^5^

Current clinical diagnosis of AKI relies primarily on changes in serum creatinine (Scr) concentration and urine output. However, these conventional indicators have limited sensitivity and often increase only after substantial renal dysfunction has already occurred, restricting early identification and timely intervention.^6^ Furthermore, despite improved understanding of SA-AKI pathophysiology, therapeutic strategies remain largely supportive, and no pharmacological intervention directly targeting the mechanisms responsible for renal injury progression is currently available. Renal hemodynamic dysregulation represents an important and potentially reversible component of SA-AKI pathogenesis. Renal blood flow (RBF) is tightly regulated by intrinsic autoregulatory mechanisms, including the myogenic response of preglomerular arterioles and tubule-glomerular feedback, which together maintain stable glomerular perfusion despite fluctuations in systemic circulation.^7,8^During sepsis, these regulatory mechanisms become impaired, resulting in heterogeneous renal microvascular perfusion characterized by regional hypoperfusion, intermittent capillary flow, and impaired oxygen delivery.^9,10^ Such disturbances may contribute to renal tissue hypoxia, inflammatory activation, and progressive tubular injury.

These observations highlight renal vascular autoregulation as a potential mechanistic link between systemic hemodynamic instability and renal injury during sepsis. Although impaired renal perfusion and microvascular dysfunction are increasingly recognized as important features of SA-AKI, the endogenous lipid signaling pathways that regulate renal vascular adaptability under septic stress remain poorly defined. In particular, whether alterations in vasoactive lipid mediators contribute to the failure of renal autoregulation during sepsis has not been fully elucidated.

20-Hydroxyeicosatetraenoic acid (20-HETE), an ω-hydroxylated metabolite of arachidonic acid generated primarily by cytochrome P450 enzymes of the CYP4A and CYP4F families, is an important endogenous regulator of vascular tone, blood pressure, and organ-specific perfusion.^11^In the kidney, 20-HETE is highly expressed within the renal vasculature and plays a critical role in regulating afferent arteriolar tone, renal autoregulation, and tubule-vascular signaling. Previous studies have shown that pharmacological or genetic inhibition of 20-HETE synthesis impairs vascular myogenic responses and disrupts renal blood flow autoregulation.^12,13^Conversely, genetic or pharmacological restoration of 20-HETE signaling protects against renal injury in experimental models by preserving renal perfusion and maintaining tubular function.^14,15^ Collectively, These findings indicate that appropriate 20-HETE activity is essential for maintaining renal vascular homeostasis. Importantly, emerging evidence suggests that 20-HETE signaling is altered during sepsis. In experimental models of septic shock, circulating 20-HETE concentrations decrease significantly and correlate with hypotension and cardiovascular dysfunction. Administration of 20-HETE analogs prevents this decline and improves vascular responsiveness, systemic blood pressure, and cardiac performance in septic animals.^16,17^ These observations raise the possibility that loss of 20-HETE signaling contributes to septic vascular dysfunction and subsequent organ injury. Mechanistically, 20-HETE has been proposed to exert its vascular effects through interaction with the G protein-coupled receptor 75 (GPR75), leading to activation of downstream signaling cascades involving phospholipase C (PLC) and protein kinase C (PKC), as well as other intracellular pathways.^18^ However, whether disruption of the 20-HETE– GPR75 signaling axis contributes to renal hemodynamic dysfunction and SA-AKI development remains unknown.

Here, we hypothesized that sepsis-induced impairment of 20-HETE signaling compromises renal vascular adaptability, thereby contributing to renal hypoperfusion and the development of SA-AKI. To test this hypothesis, we integrated longitudinal assessment of systemic and renal hemodynamics with molecular and pharmacological approaches in a CLP-induced sepsis model, complemented by a clinical cohort study in patients with sepsis. Specifically, we sought to: (1) characterize the temporal changes in circulating and renal 20-HETE during the progression of sepsis and SA-AKI, (2) determine the relationship between 20-HETE signaling and systemic and renal hemodynamic dysfunction, and (3) determine whether pharmacological restoration of 20-HETE signaling attenuates renal injury and investigate the contribution of the GPR75-associated PLC/PKC signaling pathway.

## MATERIALS AND METHODS

### Animal Preparation

All animal experiments were performed using healthy male Sprague-Dawley rats weighing 280–300g. Experimental animals were obtained from an accredited laboratory animal supplier (license number: SCXK [Beijing] 2019-0008). All procedures were approved by the Institutional Animal Care and Use Committee of the Fourth Hospital of Hebei Medical University (approval number: IACUC-4th hos Hebemu20240023) and were conducted in accordance with institutional guidelines for animal care and use. All animals were euthanized at the designated experimental endpoints following completion of the procedures.

### Experimental Model Establishment

Animals were housed under controlled environmental conditions with a 12-hour light/dark cycle, ambient temperature of 22–24°C, and relative humidity of 40–60%. Rats had free access to standard chow and water unless otherwise specified. Food was withheld for 24 hours before experimental procedures, while water remained available ad libitum.

General anesthesia was induced with inhaled isoflurane (3% for induction and 2% for maintenance). SA-AKI was induced using the CLP model to establish polymicrobial sepsis. To minimize hypothermia and maintain fluid balance during the perioperative period, warmed sterile saline (37°C, 5 mL/kg body weight) was administered subcutaneously at the dorsal neck and lower back regions immediately after surgery.

### Experimental Design and Protocols

#### Experiment 1: Temporal Characterization of Hemodynamic and 20-HETE Signaling Alterations During Sepsis

To characterize the dynamic changes associated with sepsis progression, rats were randomly assigned into six experimental groups: control group (n=6), sham-operated group (n=6), CLP-induced sepsis groups at 3, 6, 12, and 18 hours after CLP (n=6 per time point; total n=24). Sham-operated rats underwent laparotomy followed by immediate abdominal closure without manipulation of the cecum. The sepsis groups underwent CLP surgery to induce polymicrobial sepsis.

At the prespecified time points, rats underwent standardized cardiovascular and renal ultrasonographic examinations to longitudinally assess systemic cardiac function and intrarenal hemodynamics. Rats were subsequently euthanized, and blood and kidney tissues were collected for biochemical, histopathological, and molecular analyses. The experimental protocol is illustrated in Figure 1a.

**Figure 1.**
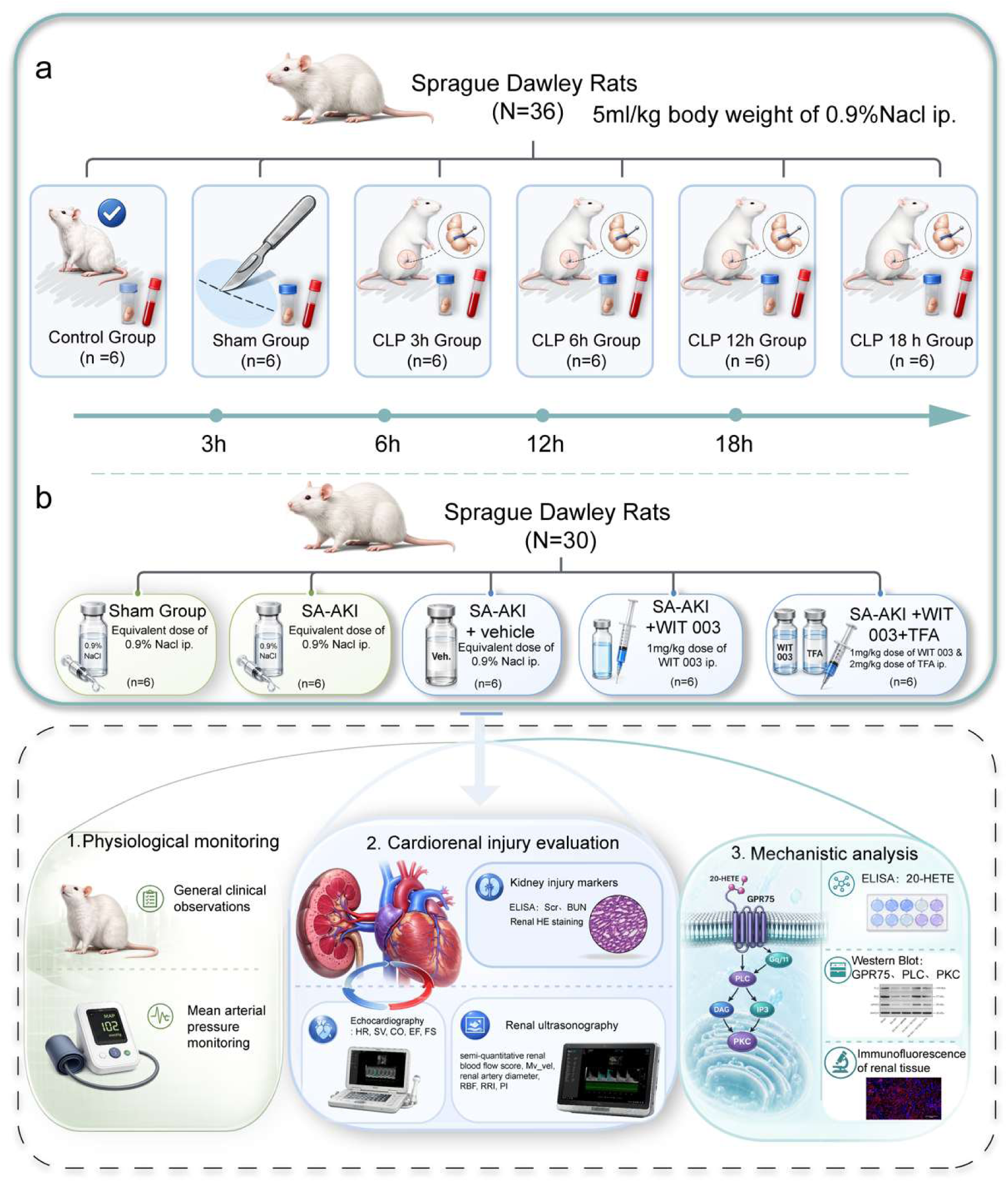
Experimental design and study protocols. (a) Experimental protocol for temporal characterization of sepsis-induced hemodynamic alterations and 20-HETE signaling changes. (b) Experimental protocol for evaluation of the therapeutic effects of WIT003 and involvement of PKC signaling in SA-AKI.

**Figure 2.**
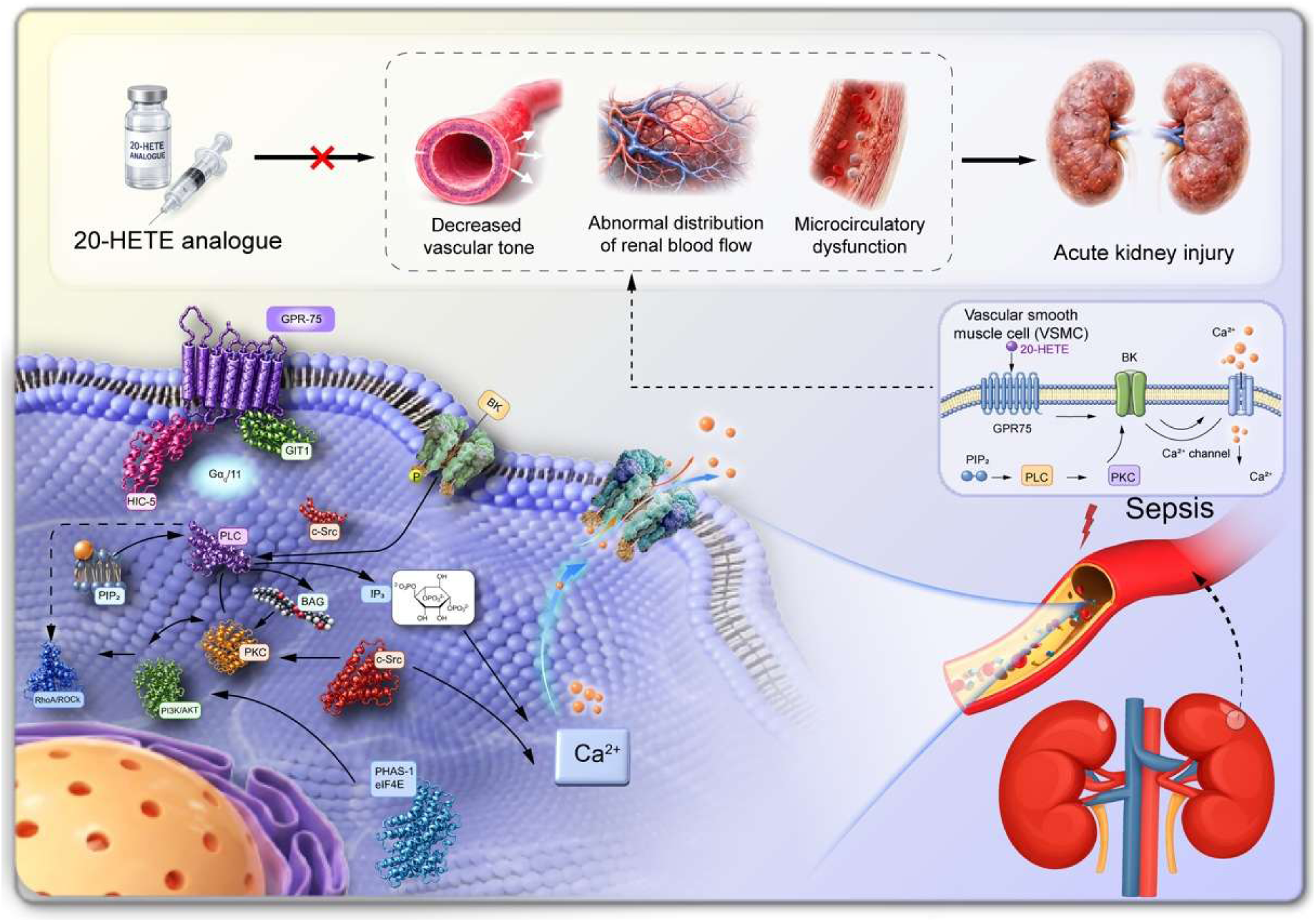
A schematic diagram illustrating the pathophysiological mechanisms of the 20-HETE– GPR75–PLC/PKC signaling pathway. In vascular smooth muscle cells (VSMCs), 20-HETE binding to GPR75 similarly activates Gαq/11 and GIT1; however, downstream effectors diverge: Gαq/11 activation stimulates protein kinase C (PKC), whereas GIT1 mediates c-Src activation. Both PKC and c-Src phosphorylate the large-conductance calcium-activated potassium (BK) channel, resulting in its functional inhibition. BK channel suppression enhances vascular tone by sensitizing the contractile apparatus to calcium. Additionally, Gαq/11 activation promotes intracellular Ca²⁺ mobilization via the phospholipase C (PLC)–inositol trisphosphate (IP₃) pathway, further contributing to 20-HETE– induced vasoconstriction. Notably, PKC activation downstream of GPR75 engagement has also been implicated as a critical mediator of 20-HETE–stimulated VSMC migration and proliferation.

#### Experiment 2: Effects of 20-HETE Restoration on SA-AKI and the Involvement of PKC Signaling

To investigate the therapeutic effects of restoring 20-HETE signaling and to determine the involvement of downstream PKC signaling in SA-AKI, rats were randomly allocated to five groups (n=6 per group): sham-operated group, SA-AKI group, SA-AKI + vehicle group, SA-AKI + 20-HETE analog group, and SA-AKI + 20-HETE analog + PKC inhibitor group.

SA-AKI was induced by CLP. The 20-HETE analog 20-5,14-HEDGE (WIT003, 1 mg/kg) and the PKC inhibitor 19–31 (TFA,2 mg/kg) were administered intraperitoneally 3 hours before CLP induction. Rats assigned to the vehicle group received an equivalent volume of the corresponding vehicle solution. In the combination-treatment group, the PKC inhibitor was administered together with WIT003 according to the same pretreatment schedule. Sham-operated and untreated SA-AKI animals underwent the corresponding procedures without administration of the active compounds.

Following CLP induction, cardiovascular and renal ultrasonography was performed according to the predefined experimental protocol. Blood and kidney tissues were subsequently collected for assessment of renal function, circulating and renal 20-HETE levels, histopathological injury, and expression of components of the GPR75–PLC/PKC signaling pathway. The experimental protocol is illustrated in Figure 1b.

#### Standardized Experimental Procedures and Sample Collection

Across both experimental protocols, animals underwent standardized assessments before and after model induction. General behavioral status was evaluated, and noninvasive blood pressure was measured using a tail-cuff system. Cardiac and renal ultrasonography were subsequently performed according to the predefined imaging protocols.

At the prespecified experimental endpoints, animals were euthanized, and blood and kidney tissues were collected for subsequent analyses. Blood samples were processed to obtain serum, which was used for measurement of Scr, blood urea nitrogen (BUN), and circulating 20-HETE concentrations according to the respective assay protocols. Kidneys were divided into two portions. One portion was fixed in paraformaldehyde, paraffin-embedded, and sectioned for hematoxylin and eosin (H&E) staining and histopathological assessment. The remaining renal tissue was rapidly snap-frozen in liquid nitrogen and stored at −80°C until subsequent Western blotting and immunofluorescence analyses.

### Ultrasound-Based Assessment of Cardiac and Renal Hemodynamics

All ultrasound examinations were performed by two independent investigators with specialized training in veterinary ultrasonography at Hebei Medical University. Operators were blinded to experimental grouping to minimize measurement bias. Animals were positioned supine and examined using a high-frequency ultrasound imaging system (Vevo 2100; VisualSonics, Toronto, Canada).

#### Cardiac Function Assessment

Cardiac function was assessed using a 2–5 MHz cardiac transducer in the parasternal short-axis view. M-mode echocardiography was used to quantify: heart rate (HR, beats/min); stroke volume (SV, μL); cardiac output (CO, mL/min); ejection fraction (EF, %); fractional shortening (FS, %).

#### Renal Ultrasound Assessment

The right kidney and renal artery were visualized using a 24-MHz high-frequency abdominal transducer positioned over the right upper abdominal region. Renal artery diameter (RAD) was measured using M-mode imaging. Renal blood flow was subsequently evaluated using color Doppler flow imaging (CDFI) and pulsed-wave Doppler techniques. Parameters included: renal blood flow score; peak systolic velocity (PSV,cm/s); end-diastolic velocity (EDV, cm/s); renal resistive index (RRI); pulsatility index (PI).

Calculation of Doppler-Derived Parameters

Mean velocity (*MV*_*_Vel_*) was estimated according to the following equation^19,20^:

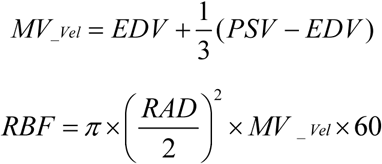

### Biochemical Analysis

Scr, BUN, and 20-HETE were quantified using commercially available assay kits according to the manufacturers’ instructions. All samples were analyzed in duplicate, and laboratory personnel performing biochemical measurements were blinded to experimental allocation.

### Renal Histological Analysis

Renal tissue sections were subjected to H&E staining. Tubular injury severity was semi-quantitatively graded using a standardized histopathological scoring system, defined as follows: Grade 0: No tubular lesions. Grade 1: Tubular lesions involving <25% of the renal cortical area. Grade 2: Tubular lesions involving 25–50% of the renal cortical area. Grade 3: Tubular lesions involving 50–75% of the renal cortical area. Grade 4: Tubular lesions involving 75–100% of the renal cortical area. For each specimen, a mean histological score was calculated from the independent evaluation of two non-overlapping tissue sections.

### Western Blot

Rat kidney tissues were homogenized in RIPA lysis buffer (Sunncell, Wuhan, China) at a fixed tissue-to-buffer ratio. Following centrifugation, the supernatant was collected and protein concentration was determined using a bicinchoninic acid assay kit (Solarbio). Equal amounts of protein were resolved by SDS-PAGE on 6–12% polyacrylamide gels and subsequently electrophoretically transferred onto polyvinylidene fluoride membranes. The membranes were blocked with 5% non-fat dry milk in Tris-buffered saline containing 0.1% Tween-20 for 1.5 to 2 h at room temperature. Thereafter, membranes were incubated overnight at 4 °C with primary antibodies against GPR75 (1:1000; Abcam), PLC (1:1000; Proteintech, Wuhan, China), PKC (1:1000; Abcam), and GAPDH (1:1000; Servicebio, Wuhan, China). After three washes with TBST, membranes were incubated with horseradish peroxidase (HRP)-conjugated secondary antibody (1:10000; Abbkine, Wuhan, China) for 1 h at room temperature. Immunoreactive bands were visualized using enhanced chemiluminescence reagent, and band intensities were quantified by densitometric analysis using ImageJ software.

### Immunofluorescence

The embedded tissue blocks were sectioned into 10-μm-thick slices and mounted onto glass slides. Primary antibodies specific to the target proteins, including PLC, PKC, and GPR75, were applied to the tissue sections and incubated at an optimized temperature. Following thorough washing to remove unbound primary antibodies, a fluorescein-labeled secondary antibody (ZF-0516; goat anti-rabbit IgG, emitting red fluorescence) was added. The immunostained sections were subsequently visualized using either a fluorescence or a confocal microscope.

### Clinical Research

#### Study design

This prospective, single-center cohort trial will conduct in the ICU of the Fourth Hospital of Hebei Medical University, planning to enroll 30 patients. The study protocol was developed by the Researcher Committee and received approval from the Ethics Committee of the Fourth Hospital of Hebei Medical University (approval number: 2025KY013). Registration for the study was completed with the China Clinical Trials Center (registration number: ChiCTR2500106881). This study adheres to the principles outlined in the *Declaration of Helsinki* (1964) and its subsequent amendments.

#### Participants

Inclusion criteria:

1. Age ≥ 18 years.
2. Diagnosis of sepsis according to the Sepsis-3 definition1 (suspected or confirmed infection, with an increase in the Sequential Organ Failure Assessment [SOFA] score of ≥ 2 points from baseline).
3. Written informed consent obtained from the patient or their legal representative.

Exclusion criteria:

1. ICU length of stay < 72 hours.
2. Pre-existing chronic kidney disease (baseline serum creatinine > 123.8 μmol/L or estimated glomerular filtration rate [eGFR] < 60 mL/min/1.73 m^2^).
3. Difficulty obtaining clear renal ultrasound images due to severe obesity (body mass index [BMI] > 40 kg/m^2^), massive ascites, severe bowel gas interference, inability to cooperate with breath-holding or position changes, or other causes.
4. Pregnancy.
5. Failure to provide signed informed consent or incomplete clinical data.

#### Data collection and ultrasound measurements

##### Data collection

Baseline demographic and clinical characteristics were collected within 24 hours of ICU enrollment. After enrollment, clinical parameters were continuously monitored and systematically recorded, including hourly measurements of body temperature, HR, respiratory rate, and MAP. Details of vasoactive medication use, including drug type, dose, and duration of administration, were also documented. For mechanically ventilated patients, ventilatory parameters, including ventilation mode, tidal volume, and oxygenation index, were recorded.

Laboratory data obtained within 6 hours of enrollment included complete blood count, renal function indices, and arterial blood gas parameters, including partial pressure of oxygen, oxygenation index, and lactate concentration. Blood samples for measurement of circulating 20-HETE concentrations were collected concurrently, processed according to a standardized protocol, and stored at −80°C until analysis. All 20-HETE measurements were performed in a single analytical batch to minimize inter-assay variability.

##### Renal POCUS

Bedside examinations were performed using a portable ultrasound system with a low-frequency convex probe. Patients were placed in the supine or semi-recumbent position and instructed to hold their breath when necessary. Standard renal views were obtained using the liver or spleen as acoustic windows.

#### Semi-quantitative scoring of renal blood flow (CDFI)

Intrarenal blood flow was evaluated using color Doppler flow imaging (CDFI) and graded semi-quantitatively on a 4-point scale according to the extent and distribution of detectable intrarenal blood flow signals:

Grade 0: No detectable blood flow signal in the kidney.

Grade 1: A scant/limited amount of blood flow signal visible at the renal hilum.

Grade 2: Evident/obvious blood flow signal at the renal hilum, with a few interlobar artery signals observed at the corticomedullary junction.

Grade 3: Blood flow signals are visible throughout the entire kidney, extending to the level of the arcuate arteries (with intact perfusion to the peripheral cortex).

##### Doppler-derived renal hemodynamic parameters

Spectral Doppler ultrasound was subsequently used to characterize renal arterial hemodynamics. Peak systolic velocity and end-diastolic velocity were measured from Doppler waveforms obtained from the renal artery or representative intrarenal arterial branches according to the predefined imaging protocol. RRI and PI were calculated as follows:

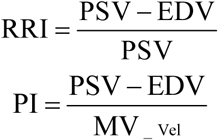

Time-averaged maximum velocity (TAMAX) and time-averaged mean velocity (TAMEAN) were also measured. For each Doppler measurement, at least 3 consecutive, technically adequate cardiac cycles were recorded. Measurements were repeated 3 times, and the mean value was used for statistical analysis. Ultrasound images and Doppler waveforms were stored digitally for subsequent offline analysis.

### Statistical Analysis

Statistical analyses were performed using IBM SPSS Statistics (version 27.0; IBM Corp., USA). Data distribution was assessed using the Shapiro–Wilk test. Normally distributed continuous variables are expressed as mean ± standard deviation (M±SD), while categorical variables are presented as numbers and percentages. For the animal experiments, comparisons among multiple experimental groups were performed using one-way ANOVA followed by Tukey’s multiple-comparison test. For time-course experiments, two-way ANOVA was used to evaluate the effects of experimental condition and time, followed by appropriate post hoc analysis. For the clinical cohort analyses, continuous variables were compared using independent-sample t tests or Mann–Whitney U tests according to data distribution.

Associations between circulating 20-HETE concentrations and renal hemodynamic parameters were evaluated using Pearson or Spearman correlation analyses as appropriate. All statistical analyses were two-sided, and a *P* value <0.05 was considered statistically significant.

## RESULTS

### Animal Experiment Part One

#### CLP-induced sepsis exhibits a biphasic systemic hemodynamic response characterized by early compensation and late cardiovascular decompensation

Following CLP induction, rats developed progressive systemic manifestations of sepsis. Beginning approximately 12 hours after surgery, CLP-treated animals exhibited characteristic signs of systemic illness, including reduced spontaneous activity, decreased food and water intake, piloerection, diminished responsiveness to external stimuli, tachypnea, tachycardia, and hypothermia. In contrast, sham-operated animals maintained normal activity and physiological status throughout the observation period.

Serial assessment of systemic hemodynamics revealed a distinct biphasic response following CLP-induced sepsis. During the early phase (≤6 hours after CLP), cardiovascular parameters remained largely preserved, with no significant differences in MAP, HR, SV, CO, EF, or FS compared with sham-operated controls, suggesting an initial compensatory hemodynamic state (Figure 3E,3F).

**Figure 3.**
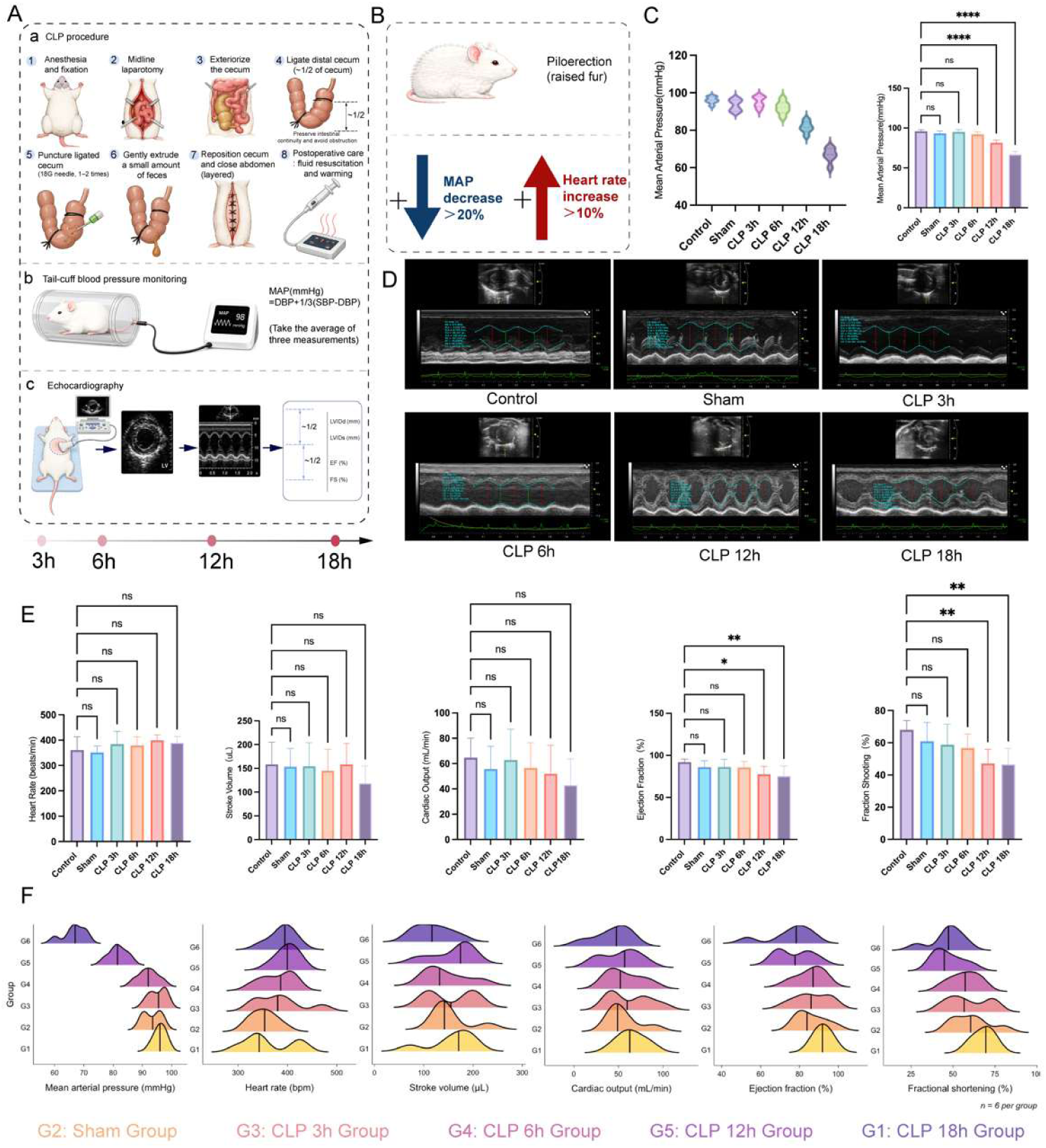
Systemic hemodynamic and ultrasound-derived parameters in rats subjected to CLP-induced sepsis. (A) Experimental workflow for hemodynamic and echocardiographic assessments. (B) Criteria for successful sepsis model establishment. (C) Hemodynamic alterations in MAP among groups. (D) Ultrasound-derived parameters among groups. (E) ultrasound-derived hemodynamic parameters among groups. (F) Distributions of systemic hemodynamic and echocardiographic parameters across groups. Data are presented as mean ± SD. \**P* < 0.05, \*\**P* < 0.01, \*\*\**P* < 0.001, **** *P* < 0.0001

By 12 hours after CLP, animals transitioned toward a decompensatory phase, characterized by a significant reduction in MAP (*P*<0.05), persistent tachycardia, and progressive impairment of cardiac performance. Although decreases in SV and CO were observed at this time point, these changes did not reach statistical significance. In contrast, left ventricular systolic function was significantly impaired, as demonstrated by reductions in EF and FS (both *P*<0.05), indicating the emergence of cardiovascular dysfunction during progressive sepsis (Figure 3E,3F).

At 18 hours after CLP, systemic hemodynamic deterioration became more pronounced. MAP declined further, while significant reductions in SV and CO were observed (*P*<0.05). EF and FS demonstrated a progressive time-dependent decline, reaching their lowest levels at the terminal observation point. Throughout the experimental period, sham-operated animals exhibited stable cardiovascular parameters without significant temporal variation.

#### Progressive renal dysfunction during sepsis is accompanied by impaired intrarenal hemodynamics and depletion of 20-HETE signaling

Renal dysfunction progressively developed following CLP-induced sepsis. Scr and BUN increased progressively compared with the control group, with the most pronounced elevations observed at 18 hours after CLP (both *P*<0.0001). At this time point, the increase in Scr was consistent with the development of clinically relevant AKI, accompanied by marked biochemical evidence of renal dysfunction (Figure 4B).

**Figure 4.**
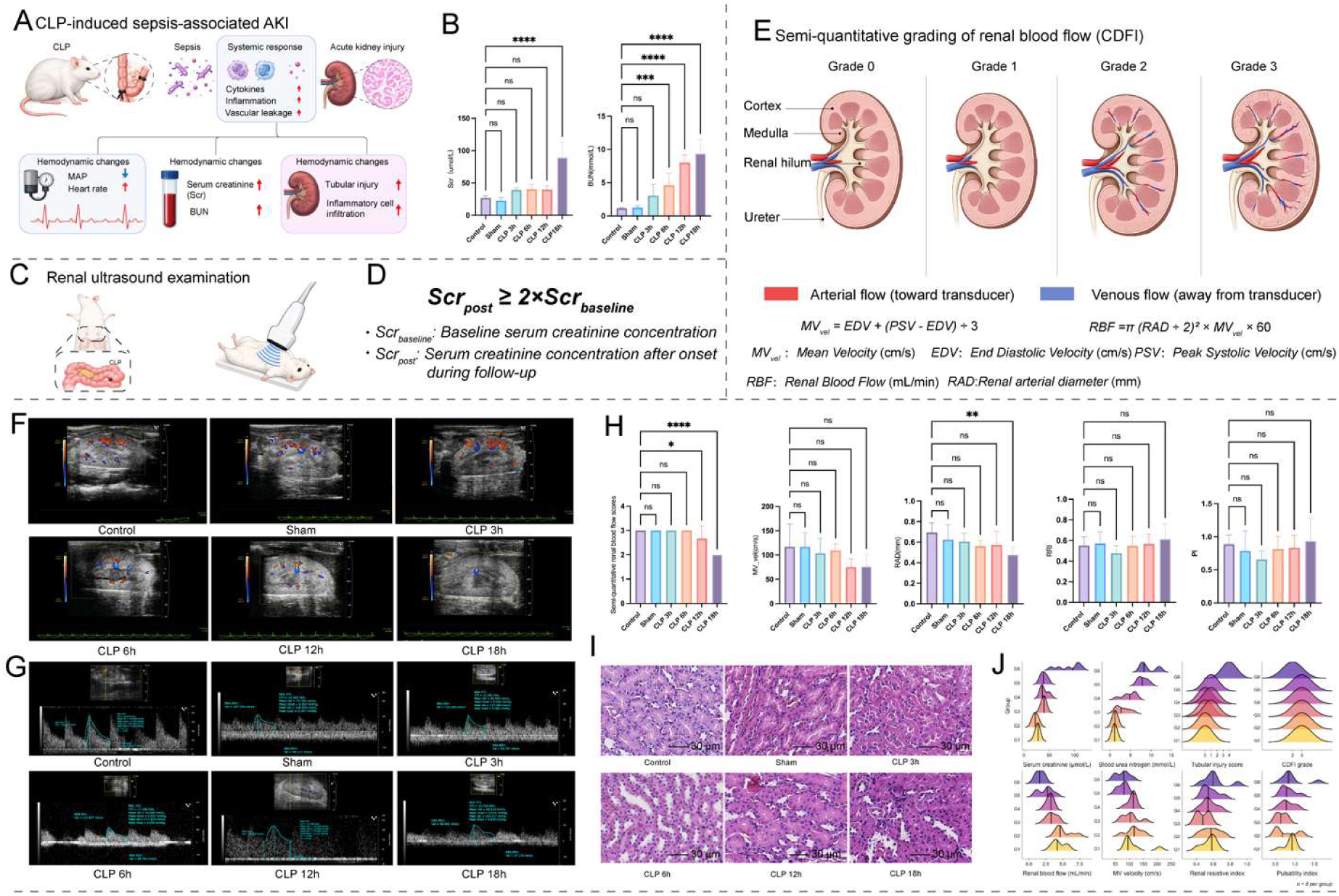
Renal function and intrarenal hemodynamic parameters in rats following CLP-induced sepsis. (A) Pathophysiological mechanisms of SA-AKI. (B) Renal functional deterioration evidenced by Scr and BUN Elevations among groups. (C) Intrarenal hemodynamic assessment by ultrasound. (D) SA-AKI modeling success criteria. (E) Semi-quantitative renal blood flow scores (CDFI) and calculation of ultrasound-derived renal hemodynamic parameters Fig.3E Semi-quantitative renal blood flow scores criteria. (F) Representative renal ultrasound images of renal semi-quantitative renal blood flow. (G) Representative renal ultrasound images. (H). Analysis of renal function and intrarenal hemodynamic parameters. (I) Tubular injury scores based on H&E-stained kidney sections. (J) Distributions of Renal function and intrarenal hemodynamic parameter across groups. All parameters were measured in the control, sham, and CLP-induced sepsis groups at the indicated time points. Data are presented as mean ± SD. \**P* < 0.05, \*\**P* < 0.01, \*\*\**P* < 0.001, **** *P* < 0.0001

Histopathological analysis further demonstrated progressive structural injury in the kidneys of CLP-treated animals. H&E staining revealed interstitial edema, vacuolar degeneration of tubular epithelial cells, loss of the tubular brush border, and tubular luminal obstruction. The severity of these lesions increased progressively with the duration of sepsis, with the highest tubular injury scores observed at 18 hours after CLP (*P*<0.05). No significant differences in renal histopathological findings were observed between control and sham-operated animals (Figure 4I, 4J).

Renal ultrasonography demonstrated a parallel deterioration in intrarenal hemodynamics during sepsis Semiquantitative assessment of renal blood flow revealed a progressive reduction beginning at 12 hours after CLP, reaching the lowest level at 18 hours (*P*<0.05). Mean renal arterial velocity (MV_*_Vel_*) also declined progressively, accompanied by a reduction in RAD, with both parameters showing the greatest changes at 18 hours (*P*<0.05). Consistent with these alterations, estimated RBF decreased progressively during sepsis and was significantly reduced at 18 hours compared with controls (*P*<0.05). RRI and PI showed a gradual upward trend during disease progression, although neither reached statistical significances (Figure 4F, 4G, 4H).

#### Early loss of 20-HETE signaling accompanies progressive renal hemodynamic dysfunction during sepsis

We next examined whether alterations in renal hemodynamics were associated with changes in 20-HETE signaling. Circulating 20-HETE concentrations remained unchanged in sham-operated animals compared with controls. In contrast, both circulating and renal 20-HETE levels decreased as early as 3 hours after CLP and progressively declined thereafter, reaching their lowest levels at 18 hours. This reduction preceded the most pronounced deterioration in renal function and renal blood flow, suggesting that depletion of 20-HETE signaling may represent an early event during the progression of septic renal dysfunction (Figure 5 II).

**Figure 5.**
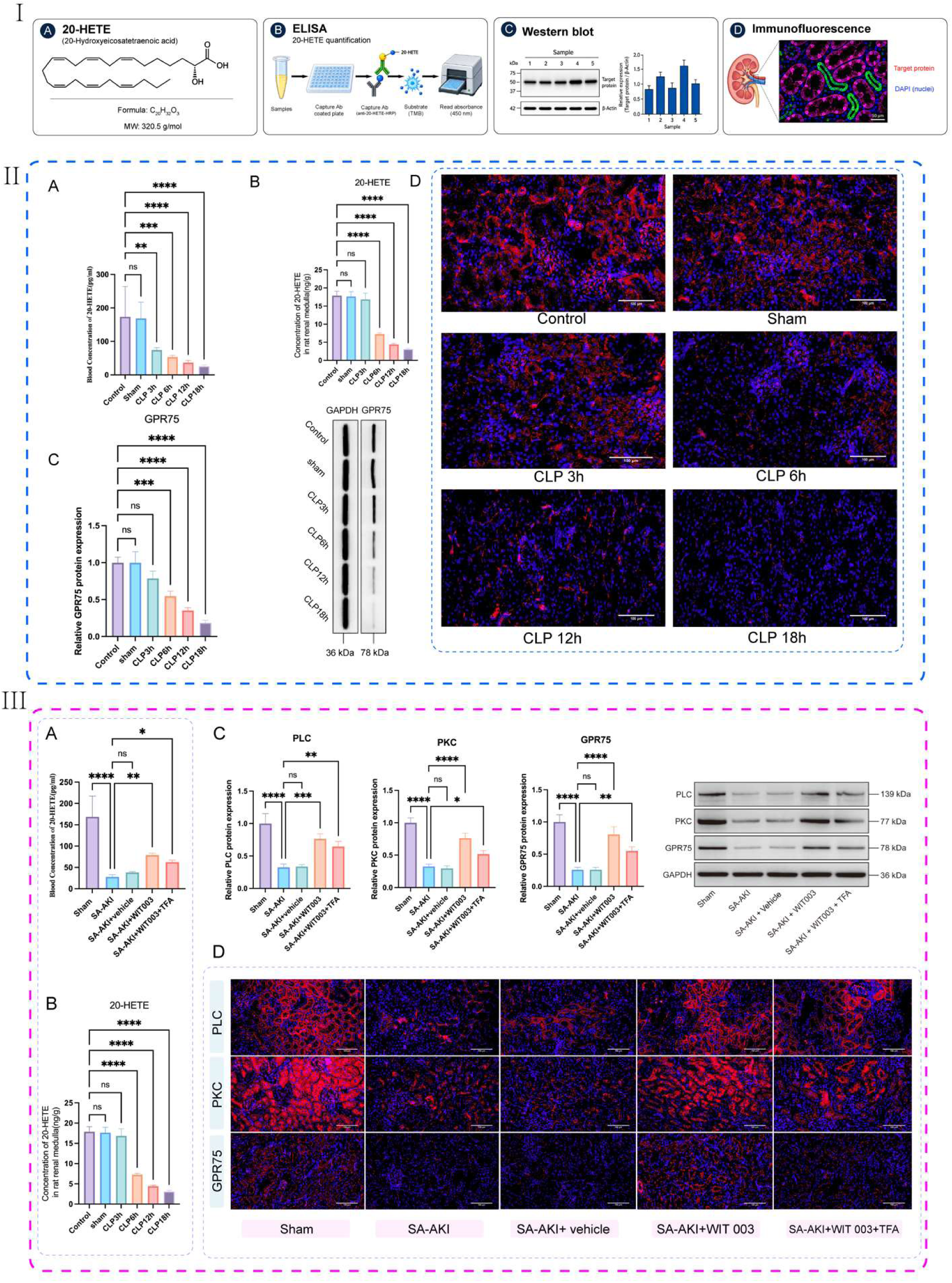
Blood 20-HETE levels and renal 20-HETE and GPR75 expression in a rat model of sepsis and SA-AKI. ((I) Experimental methods for blood and tissue specimens. (II) Blood 20-HETE levels and renal 20-HETE and GPR75 expression in a rat model of sepsis: (A) Time course of blood 20-HETE levels in rats subjected to CLP-induced sepsis. (B) Time course of renal 20-HETE levels in rats subjected to CLP-induced sepsis. (C) Representative Western blot images and quantitative analysis of GPR75 protein expression in kidney tissues. (D) Representative immunofluorescence images showing GPR75 expression in kidney tissues across groups. (III) 20-HETE levels and renal 20-HETE-GPR75-PLC/PKC axis protein expression in SA-AKI rat model: (A) Blood 20-HETE levels in different groups. (B) Renal tissue 20-HETE levels in different groups. (C) Representative Western blot images and quantitative analysis of 20-HETE-GPR75-PLC/PKC axis protein expression in kidney tissues. (D) Representative immunofluorescence images showing 20-HETE-GPR75-PLC/PKC axis expression in kidney tissues across experimental. Data are presented as mean ± SD. \**P* < 0.05, \*\**P* < 0.01, \*\*\**P* < 0.001, \*\*\*\**P* < 0.0001

Consistent with the reduction in 20-HETE availability, renal expression of G protein-coupled receptor 75 (GPR75), a proposed receptor for 20-HETE, also decreased progressively during sepsis. Immunofluorescence analysis further demonstrated alterations in the renal distribution and abundance of GPR75 and its downstream signaling components across the experimental groups (Figure 5 II). Together, these findings establish a temporal association among depletion of 20-HETE signaling, impaired intrarenal hemodynamics, and progressive renal injury during CLP-induced sepsis.

### Animal Experiment Part Two

#### 20-HETE Analog Administration Improves Systemic Hemodynamics in SA-AKI

To determine whether restoration of 20-HETE signaling could ameliorate the hemodynamic abnormalities associated with SA-AKI, we next evaluated systemic cardiovascular parameters following administration of the 20-HETE analog 20-5,14-HEDGE (WIT003).

Compared with untreated SA-AKI rats, administration of WIT003 significantly increased MAP (*P*<0.05), indicating improved systemic arterial pressure during established SA-AKI (Figure 6 I). In contrast, co-administration of WIT003 with the TFA resulted in only a modest increase in MAP compared with the SA-AKI group, which did not reach statistical significance. Moreover, the increase in MAP was attenuated relative to that observed with WIT003 treatment alone. No significant difference in MAP was observed between the SA-AKI and SA-AKI + vehicle groups, confirming that the vehicle itself did not materially affect systemic hemodynamics (Figure 6 I).

**Figure 6.**
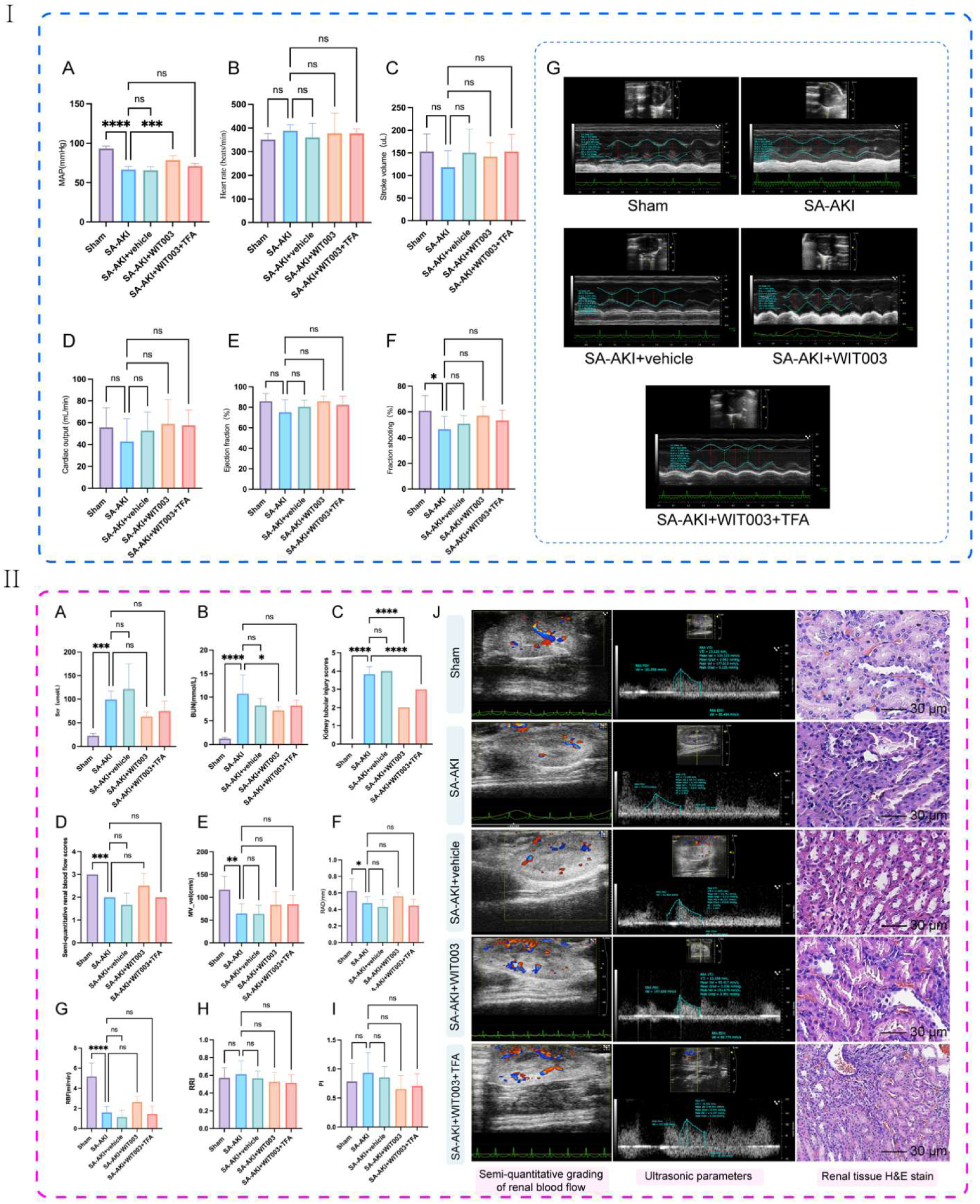
Systemic hemodynamic and ultrasound-derived parameters & changes in renal function and intrarenal hemodynamics in a rat model of CLP-induced SA-AKI. (I) Systemic hemodynamic and ultrasound-derived parameters in a rat model of CLP-induced SA-AKI: (A) Mean arterial pressure. (B) Heart rate. (C) Stroke volume. (D) Cardiac output. (E)Ejection fraction. (F)Fractional shortening. (G) Cardiac ultrasound-derived parameters. Data are presented as mean ± SD (n = 6 per group per time point). (II) The changes in renal function and intrarenal hemodynamics in a rat model of SA-AKI: (A) Scr levels. (B) BUN levels. (C) Tubular injury scores based on H&E-stained kidney sections. (D)Semi-quantitative renal blood flow scores. (E)Mean flow velocity (MV__Vel_). (F) Renal artery luminal diameter. (G) Estimated renal blood flow. (H) Renal resistive index. (I) Pulsatility index. (J) Representative renal ultrasound images and images of H&E staining of renal tissue in the sham, SA-AKI, SA-AKI+vehicle, S-AKI+WIT003, SA-AKI+WIT003+TFA groups at the indicated time points. Data are presented as mean ± SD. \**P* < 0.05, \*\**P* < 0.01, \*\*\**P* < 0.001, \*\*\*\**P* < 0.0001

HR remained comparable among all experimental groups (Figure 6 I B). WIT003-treated rats also exhibited numerical increases in SV and CO compared with untreated SA-AKI rats, although these differences did not reach statistical significance (Figure 6 I C,6 I D). Notably, the magnitude of these changes was greater following WIT003 monotherapy than following combined WIT003 and TFA treatment. A similar pattern was observed for indices of left ventricular systolic function, including EF and FS (Figure 6 I E,6 I F). However, neither parameter differed significantly among the experimental groups.

Collectively, these findings demonstrate that restoration of 20-HETE signaling with WIT003 significantly improves systemic arterial pressure in SA-AKI, whereas concomitant TFA attenuates this hemodynamic response. Although changes in CO and systolic function did not reach statistical significance, their directional patterns were consistent with a potential improvement in cardiovascular performance following WIT003 administration.

#### 20-HETE Restoration Attenuates Renal Injury and Improves Intrarenal Hemodynamics in SA-AKI

Having demonstrated that WIT003 improved systemic arterial pressure in SA-AKI, we next examined whether restoration of 20-HETE signaling could ameliorate renal dysfunction and intrarenal hemodynamic abnormalities.

Compared with untreated SA-AKI rats, WIT003 administration reduced Scr and BUN levels (Figure 6 II A,6 II B), accompanied by a significant improvement in renal histopathological injury (*P*<0.05) (Figure 6 II C). Histological analysis further demonstrated attenuation of tubular structural abnormalities following WIT003 treatment. Although combined administration of WIT003 and the PKC inhibitor also resulted in improvements in renal biochemical and histopathological parameters compared with the untreated SA-AKI group, these protective effects were less pronounced than those observed with WIT003 alone (Figure 6 II C, 6 II J).

We next assessed whether the reno-protective effects of WIT003 were accompanied by improvement in intrarenal hemodynamics. WIT003 treatment did not significantly alter RAD (Figure 6 II F) but increased mean renal arterial velocity (MV_*Vel*) (Figure 6 II E) and estimated RBF (Figure 6 II G). In parallel, RRI (Figure 6 II H) and PI (Figure 6 II I) were reduced following WIT003 administration, consistent with improved renal vascular perfusion and hemodynamic adaptation.

In animals receiving combined WIT003 and TFA treatment, the improvements in renal perfusion parameters were attenuated compared with those observed with WIT003 alone. Nevertheless, these parameters remained directionally improved relative to untreated SA-AKI animals. The differences between the combination-treatment and SA-AKI groups did not reach statistical significance.

#### 20-HETE Restoration Reconstitutes the Renal GPR75–PLC/PKC Signaling Axis in SA-AKI

We next investigated whether the protective effects of WIT003 were associated with restoration of the renal 20-HETE–GPR75–PLC/PKC signaling axis. Compared with untreated SA-AKI rats, administration of WIT003 significantly increased circulating 20-HETE concentrations and was accompanied by increased renal protein expression of GPR75, PLC, and PKC (*P*<0.05) (Figure 5 III). To further assess the involvement of PKC signaling, WIT003 was administered in combination with TFA. Although 20-HETE concentrations and the expression levels of GPR75, PLC, and PKC remained directionally higher in the combination-treatment group than in untreated SA-AKI animals, the magnitude of these changes was attenuated compared with WIT003 monotherapy (Figure 5 III). These findings indicate that restoration of 20-HETE availability is associated with reactivation of the renal GPR75–PLC/PKC signaling axis and further support the involvement of PKC signaling in the renal protective effects of 20-HETE during SA-AKI.

### Clinical Research Part

#### Circulating 20-HETE Levels Are Markedly Reduced in Sepsis and Further Depleted in Patients With SA-AKI

A total of 30 participants were enrolled in the clinical cohort, including 10 healthy adults, 10 patients with sepsis without AKI, and 10 patients with SA-AKI.

Circulating 20-HETE concentrations differed markedly among the 3 groups. Plasma 20-HETE levels were substantially lower in patients with sepsis without AKI than in healthy adults (18.63±1.57 versus 221.74±35.62 pg/mL), and were further reduced in patients with SA-AKI (12.55±3.49 pg/mL, *p*<0.001 across groups). Thus, circulating 20-HETE levels demonstrated a graded reduction from health to sepsis and further to SA-AKI, suggesting that progressive depletion of 20-HETE is associated with the development of renal dysfunction in sepsis.

We next examined whether changes in circulating 20-HETE were accompanied by alterations in renal Doppler-derived hemodynamic parameters. Compared with septic patients without AKI, patients with SA-AKI exhibited a significantly lower PI [1.07(0.40) versus 1.27(0.23); *P*=0.002]. MV_*_Vel_* was also significantly reduced in the SA-AKI group compared with septic patients without AKI (18.39(5.72) versus 21.46(3.07) cm/s, *P*=0.007). These findings indicate that SA-AKI was associated with reduced renal blood-flow velocity and altered renal arterial pulsatility (See Sup.Table 1).

In contrast, EDV, RRI, and TAMEAN did not differ significantly among the clinical groups. The differential behavior of these Doppler-derived parameters suggests that conventional renal vascular indices may not uniformly capture the hemodynamic alterations associated with SA-AKI. Importantly, the marked reduction in circulating 20-HETE was accompanied by a decrease in MV__Vel_ in patients with SA-AKI, linking impaired 20-HETE availability with altered renal hemodynamics in the clinical setting. Together with the findings from the experimental model, these observations support an association between depletion of 20-HETE signaling and impaired renal perfusion during sepsis and provide clinical evidence for the translational relevance of the 20-HETE–renal hemodynamic axis.

## DICUSSION

In the present study, we identified a previously underappreciated link between impaired 20-HETE signaling and renal hemodynamic dysfunction during SA-AKI. Using a temporally resolved CLP model, we found that sepsis evolved from an early compensatory cardiovascular state to a late decompensated state characterized by hypotension, impaired cardiac performance, progressive renal dysfunction, and deterioration of intrarenal blood flow. Importantly, circulating and renal 20-HETE levels declined early during sepsis and reached their lowest levels at the time of maximal renal and systemic hemodynamic disturbance. This temporal relationship was accompanied by downregulation of renal GPR75 and downstream PLC/PKC signaling.

Restoration of 20-HETE signaling with WIT003 improved systemic arterial pressure, renal hemodynamics, renal biochemical indices, and tubular histopathology in septic rats. These effects were attenuated by pharmacological inhibition of PKC, providing pharmacological evidence that PKC-associated signaling contributes to the protective effects of 20-HETE restoration. Finally, our clinical observations demonstrated a marked reduction in circulating 20-HETE concentrations in patients with sepsis, with a further decrease in patients with SA-AKI, accompanied by reduced renal blood-flow velocity and altered renal Doppler parameters. Together, these findings support a model in which loss of 20-HETE signaling contributes to impaired renal vascular adaptation during sepsis and thereby facilitates the development of SA-AKI.

### 1. Sepsis-associated renal dysfunction is temporally linked to progressive cardiovascular and intrarenal hemodynamic failure

A major observation from our study is the biphasic nature of the systemic hemodynamic response following CLP. During the early phase of sepsis, systemic blood pressure and cardiac function were relatively preserved, whereas progressive cardiovascular deterioration became evident at later time points. By 12 to 18 hours after CLP, MAP, SV, and CO were reduced, accompanied by deterioration in EF and FS. In parallel, RBF progressively declined and renal injury became more pronounced.

This observation is consistent with the increasingly recognized concept that SA-AKI cannot be explained simply by a sustained reduction in global renal blood flow. Human and experimental studies have demonstrated that SA-AKI may develop in the setting of preserved or even hyperdynamic systemic circulation, while regional and microvascular perfusion becomes heterogeneous.^21^ The renal microcirculation is particularly vulnerable because renal oxygen delivery and utilization depend on a highly specialized vascular architecture, and sepsis can induce heterogeneous perfusion, endothelial dysfunction, altered oxygen extraction, glycocalyx disruption, and inflammatory activation.^22^

Our temporal observations extend these concepts by demonstrating that systemic cardiovascular decompensation and deterioration of intrarenal hemodynamics evolve in parallel with depletion of 20-HETE. The 18-hour time point was therefore selected for mechanistic intervention because it represented the convergence of maximal renal dysfunction, cardiovascular deterioration, impaired renal perfusion, and suppression of the 20-HETE/GPR75 axis. This temporal resolution is important because it places 20-HETE depletion within the evolving pathophysiological sequence rather than viewing it simply as a late consequence of established kidney injury.

The current consensus recognizes SA-AKI as a heterogeneous syndrome defined by sepsis together with KDIGO-defined AKI occurring within the sepsis time window.^5^Recent clinical data further demonstrate that SA-AKI is common in the ICU and is associated with substantial morbidity and mortality.^23^ Thus, identifying molecular processes that occur during the transition from systemic compensation to renal dysfunction may be particularly important for therapeutic development.

### 2. Our findings identify 20-HETE depletion as a potential determinant of impaired renal vascular adaptation

20-HETE is a major CYP4A/CYP4F-derived metabolite of arachidonic acid and is highly expressed in the kidney and vascular system. It has traditionally been characterized as a potent vasoconstrictor and an important mediator of vascular myogenic responses, tubule-glomerular feedback, and renal blood-flow autoregulation.^24^

This biological function provides an important framework for interpreting our findings. Although 20-HETE is commonly viewed as a vasoconstrictor, the physiological role of a vasoconstrictor mediator cannot be equated simply with pathological vasoconstriction. Appropriate dynamic regulation of vascular tone is essential for autoregulation. In the renal circulation, 20-HETE contributes to the ability of preglomerular arterioles to respond to changes in perfusion pressure. Experimental inhibition of 20-HETE synthesis impairs myogenic responses and renal blood-flow autoregulation.^25^

Our observation that 20-HETE levels decreased early after CLP, before the most severe biochemical and histological renal injury, therefore raises the possibility that loss of appropriate 20-HETE signaling may impair renal vascular adaptation rather than simply causing excessive vasoconstriction.

This interpretation also helps reconcile apparently conflicting literature regarding 20-HETE and kidney injury. In some ischemia-reperfusion models, inhibition of 20-HETE synthesis or antagonism of 20-HETE signaling was protective, whereas other studies demonstrated that 20-HETE analogs protected against renal ischemia-reperfusion injury by preserving medullary blood flow and reducing tubular sodium transport.^26^These apparently divergent effects likely reflect the highly context-dependent biology of 20-HETE, including differences in disease model, timing, tissue compartment, magnitude of 20-HETE production, and balance between vascular and tubular effects.

Our findings therefore support a more nuanced model in which the biological consequence of 20-HETE signaling depends on whether the system is experiencing excess, deficiency, or inappropriate spatial/temporal regulation of the mediator.

### 3. Restoration of 20-HETE signaling improves both systemic and renal hemodynamics

The therapeutic experiments provide functional support for this hypothesis. Administration of WIT003 significantly increased MAP in septic rats and was accompanied by directional improvements in SV and CO. More importantly, WIT003 improved renal perfusion, with increased MV__Vel_ and estimated RBF and reductions in RRI and PI, together with attenuation of biochemical and histological renal injury.

These findings are consistent with previous experimental studies showing that 20-HETE analogs can improve vascular responsiveness during endotoxemia. In rodent models of septic shock, 5,14-HEDGE has been reported to prevent vascular hypo-reactivity, hypotension, and tachycardia and to modulate nitric oxide, prostanoid, and inflammatory pathways.^14^

Our study extends these observations in several important ways.First, previous studies largely focused on systemic vascular reactivity during endotoxemic shock, whereas our CLP model addresses polymicrobial sepsis and incorporates serial assessment of renal hemodynamics. Second, our study links restoration of 20-HETE signaling not only to recovery of MAP but also to improvement in intrarenal blood-flow characteristics. Third, these hemodynamic effects were accompanied by attenuation of tubular injury, suggesting that restoration of vascular adaptation may contribute to preservation of renal tissue integrity. The latter point is particularly relevant because SA-AKI is increasingly understood as a disorder involving interactions between systemic hemodynamics, regional perfusion, microvascular dysfunction, inflammation, and tubular metabolic adaptation rather than as a simple ischemic tubular necrosis syndrome.^27^

### 4. The 20-HETE–GPR75–PLC/PKC axis provides a mechanistic link between lipid signaling and vascular function

A central mechanistic finding of this study is the coordinated reduction of 20-HETE and GPR75, accompanied by suppression of PLC/PKC signaling during progressive sepsis. Conversely, WIT003 restored 20-HETE concentrations and increased renal GPR75, PLC, and PKC protein abundance.

The identification of GPR75 as a receptor for 20-HETE provides an important molecular framework for these observations.^28^ Garcia et al. demonstrated that 20-HETE interacts with GPR75 and activates Gαq/11-dependent signaling, while GIT1-dependent pathways engage PKC and c-Src signaling in vascular cells. In vascular smooth muscle cells, these pathways influence BK channel phosphorylation and vascular contractility.^29^Our findings are therefore mechanistically concordant with established vascular biology while extending it into the context of SA-AKI. Importantly, PKC inhibition partially attenuated the effects of WIT003 on MAP, renal function, and renal hemodynamic parameters. This pharmacological interaction provides evidence that PKC-associated signaling contributes to the protective phenotype. However, these findings should not be interpreted as definitive proof that PKC is the sole downstream effector of GPR75. GPR75 can engage multiple signaling branches, including Gαq/11–PLC/IP3/PKC and GIT1/c-Src/EGFR pathways.^30^Accordingly, our data support a model in which 20-HETE restoration reactivates GPR75-associated signaling, with PKC representing an important—but probably not exclusive—downstream mediator of vascular and renal protection.This distinction will be important for future mechanistic studies.

### 5. Why does loss of a vasoconstrictor cause renal hypoperfusion?

An apparent paradox emerging from our study is that reduction of a traditionally vasoconstrictive mediator is associated with reduced renal blood flow rather than renal vasodilation with preserved perfusion. We propose that this apparent contradiction reflects the physiological role of 20-HETE in vascular autoregulation rather than absolute vascular resistance. Under physiological conditions, the renal circulation must continuously adjust vascular resistance to maintain relatively stable renal blood flow despite fluctuations in systemic pressure. 20-HETE participates in the myogenic response of preglomerular arterioles and in tubule-glomerular feedback.^31^During sepsis, the problem may therefore not be simply “too much vasoconstriction” or “too little vasoconstriction,” but rather loss of appropriate vascular responsiveness

### 6. The clinical data provide translational support for a 20-HETE–renal hemodynamic phenotype

The human cohort provides an important translational dimension to the experimental findings. Circulating 20-HETE concentrations were dramatically reduced in patients with sepsis and were further reduced in patients with SA-AKI. This stepwise pattern parallels the temporal depletion observed in the animal model.

Importantly, reduced 20-HETE levels were accompanied by reduced MV_*_Vel_* and altered PI in patients with SA-AKI. These findings suggest that 20-HETE depletion is associated with measurable abnormalities in renal vascular hemodynamics in humans. The reduction in PI is particularly interesting. RRI is widely used to assess renal vascular impedance, but its interpretation in critically ill patients is complex because it is influenced by systemic arterial pressure, heart rate, arterial compliance, venous pressure, and cardiac function. Consequently, a normal or modestly altered RRI does not necessarily exclude clinically relevant renal microcirculatory dysfunction.

However, we deliberately avoid labeling this phenotype definitively as “vasomotor paralysis” or “cortical hypoperfusion,” because renal Doppler ultrasound does not directly measure cortical microvascular flow or tissue oxygenation. Rather, our data support the more conservative concept of an altered renal hemodynamic phenotype associated with reduced 20-HETE availability. This distinction will be important for subsequent clinical validation.

### Limitations

Several limitations should be acknowledged. Firstly, our mechanistic inference relies primarily on pharmacological intervention. Although PKC inhibition attenuated the beneficial effects of WIT003, pharmacological inhibitors may have off-target effects. Genetic approaches, including GPR75 loss-of-function and PKC isoform-specific manipulation, will be necessary to establish causal relationships. Secondly, the specific PKC isoform responsible for the observed phenotype remains unknown. Previous vascular studies have implicated PKCα in 20-HETE/GPR75 signaling, but our experiments did not resolve isoform-specific contributions. Thirdly, our renal ultrasound measurements provide an estimate of renal blood flow and vascular impedance but do not directly quantify renal cortical or medullary microcirculation. Advanced techniques such as contrast-enhanced ultrasound, laser speckle imaging, intravital microscopy, renal tissue oxygenation measurements, or MRI-based perfusion imaging could provide complementary evidence.

### Conclusions

In summary, these findings support a model in which loss of the 20-HETE–GPR75–PLC/PKC signaling axis contributes to impaired renal vascular adaptation during sepsis and thereby promotes SA-AKI. Rather than viewing 20-HETE solely as a vasoconstrictor mediator, our findings suggest that its physiological importance may lie in maintaining appropriate vascular responsiveness and autoregulatory capacity during hemodynamic stress. Future studies integrating GPR75 genetic manipulation, PKC isoform-specific approaches, direct assessment of renal microcirculation and tissue oxygenation, and larger prospective clinical cohorts will be necessary to establish the causal and translational significance of this pathway.

## Grants

Funding Declaration:

Natural Science Foundation of Hebei Province, China (No: H2023206071) Medical Science Research Project of Hebei, China (No: 20260518)

## Author Contributions

YQ.Han and ZJ.Hu conceived and designed the study. YQ.Han and LY.Li developed the study protocols. HX. Zhang, RK. Yang, and Xin Lin conducted the animal study and performed the molecular and biological assays. LX. Liu and YH.Chen executed the mass spectrometric analysis. YL.Yin acquired the clinical data. CC. Zhao and YQ.Han performed the pathological evaluations. YQ. Han and LY. Li conducted the statistical analyses. YQ.Han interpreted the data and drafted the manuscript.ZJ.Hu supervised the project and verified the integrity and accuracy of the study. All authors reviewed, edited, and approved the final manuscript.

---

**Supplemental Table 1.** Comparison of serum 20-HETE levels and renal ultrasound parameters of enrolled participants.

| Indictors | Healthy Adults<br>(n=10) | Control Group<br>(n=10) | SA-AKI Group<br>(n=10) | *P* Value |
| --- | --- | --- | --- | --- |
| 20-HETE<br>(pg/mL) | 221.74±35.62 | 18.63±1.57 | 12.55±3.49 | <0.001 |
| PSV (cm/s) | 36.75±3.18 | 40.74±11.38 | 31.36±6.86 | 0.053 |
| EDV (cm/s) | 14.59(2.11) | 12.14(5.57) | 11.25(4.76) | 0.152 |
| MV<sub>vel</sub> (cm/s) | 21.46(3.07) | 20.47(9.83) | 18.39(5.72) | 0.007 |
| RRI | 0.57(0.04) | 0.71(0.08) | 0.625(0.15) | 0.147 |
| PI | 1.24(0.08) | 1.27(0.23) | 1.07(0.4) | 0.002 |
| TAMEAN (cm/s) | 12.56(0.84) | 8.90(4.89) | 7.79(2.76) | 0.101 |
| TAMAX (cm/s) | 22.82±1.08 | 18.73±5.41 | 17.56±6.22 | 0.052 |

